# Functional Difference of Mitochondrial Genome and Its Association with Traits of Common Complex Diseases in Humans

**DOI:** 10.1101/189845

**Authors:** Young Min Cho, Kyong Soo Park, Youngmi Kim Pak, Masashi Tanaka, Hong Kyu Lee

**Author notes:** **Corresponding author**: Hong Kyu Lee, MD, PhD, Department of Internal Medicine, College of MedicineEulji UniversitySeoulSouth Korea.

## Abstract

Recent evidence suggests that mitochondrial genomes harboring common mitochondrial DNA polymorphisms might have functional difference and could be associated with common complex human diseases such as metabolic syndrome and cancer that are related to mitochondrial dysfunction. However, there has been no report examining the functional difference of mitochondrial genome in the pathogenesis of such diseases at the cellular or molecular level. In order to examine the effect of mitochondrial genome on metabolic syndrome or cancer without interference from nuclear genes, we analyzed *trans*-mitochondrial cytoplasmic hybrid cells (cybrids) with common Asian mtDNA haplogroups A, B, D, and F from healthy volunteers. The mitochondrial oxygen consumption rates of cybrids were associated with multiple components of metabolic syndrome such as body mass index, waist circumference, serum triglyceride levels and high-density lipoprotein cholesterol levels. In addition, the cybrids showed varying degree of tumorigenicity both *in vitro* and *in vivo*. Especially, the cybrids harboring mtDNA haplogroup D had a significantly slower growth rate. These findings suggest that the phenotypes of common complex diseases in humans can be determined by their mitochondrial genomes. Therefore, not only nuclear genome but also mitochondrial genome should be considered in explaining the genetic pathogenesis of common complex human diseases.

## Introduction

Mitochondria are the intracellular organelles that produce most of the energy needed to maintain life through the process of oxidative phosphorylation (OXPHOS). They contain their own DNA (mitochondrial DNA; mtDNA) encoding 13 subunits of electron transfer chains and 2 rRNAs and 22 tRNAs for their protein synthesis machinery (Poyton and McEwen 1996; Wallace 2005). The human mtDNA shows extensive polymorphisms, which have long been considered to be neutral and have no advantage or disadvantage (Kimura 1968). However, there is a growing body of evidence challenging this idea. Genetic association studies have revealed that the common human mtDNA variants may influence the longevity (De Benedictis et al. 1999; Niemi et al. 2003; Tanaka et al. 1998), type 2 diabetes mellitus (Fuku et al. 2007), or the penetrance of certain mitochondrial diseases (Brown et al. 2002; Howell et al. 2003). It was suggested that the geographic region-specific distribution of mtDNA haplogroups (specific lineages of related mtDNA haplotypes) were formed by natural selection against the influence of latitude or thermal environments (Mishmar et al. 2003; Ruiz-Pesini et al. 2004). In accordance with this idea, common mtDNA polymorphisms or mitochondrial genomes were shown to alter mitochondrial matrix pH and intracellular calcium dynamics in human cells (Kazuno et al. 2006) and contribute to the production of different reactive oxygen species (ROS) in murine cells (Moreno-Loshuertos et al. 2006). It was also suggested that mitochondrial genome might contribute to the pathogenesis of common complex human diseases such as cancer (Brandon et al. 2006) or metabolic syndrome (a cluster of abnormalities such as obesity, hyperglycemia, hypertriglyceridemia and low levels of serum high density lipoprotein [HDL] cholesterol) that are related to mitochondrial dysfunction (Wilson et al. 2004). However, the phenotypic effects of variation in the mitochondrial genome have been considered to be difficult to determine due to the variations in the nuclear genome, epigenetic effect such as imprinting, and multiple environmental factors. It was very recently shown that conplastic strains of rats with identical nuclear background but different mitochondrial genome harboring multiple naturally occurring polymorphisms demonstrate distinct metabolic phenotypes regarding to glucose homeostasis (Pravenec et al. 2007). In our current study, to examine the effect of mitochondrial genome on metabolic syndrome or cancer without interference from nuclear genome, we examined the characteristics of *trans*-mitochondrial cytoplasmic hybrid cells (cybrids), which share identical nuclear genome but have different mitochondrial genome (King and Attardi 1989).

## Results

We introduced mitochondrial genomes harboring common Asian-specific haplogroups A, B, D and F (n = 6 in each group) (Wallace et al. 1999) into an mtDNA-less osteosarcoma cell line (143B TK^‒^ ρ^0^) by the cybrid technique using platelets from volunteers who were free from clinically diagnosed cancer at the time of this study. Among the 24 cybrid clones, 3 clones (cybrid F1, F2 and F3) were excluded from analysis for the metabolic syndrome, since the mtDNA donors of these clones had type 2 diabetes, which might cause secondary mitochondrial dysfunction (Kelley et al. 2002). The remaining 21 subjects had neither metabolic syndrome nor diabetes.

### Associations with metabolic syndrome

Firstly, we examined whether the respiratory function of the cybrids might be associated with the phenotypes of metabolic syndrome in the mtDNA donors (n = 21). The basal oxygen consumption rate (OCR) and non-mitochondrial OCR after KCN treatment were measured for each cybrid. We calculated the mitochondrial OCR by subtracting the non-mitochondrial OCR from the basal OCR and compared it with the metabolic phenotypes of the mtDNA donors.

The mitochondrial OCR showed a good correlation with body mass index (BMI) (r = ‒0.4541, *P* < 0.05), waist circumference, hip circumference, serum triglyceride level and HDL cholesterol level (Fig. 1). Although it was not correlated with insulin resistance index assessed by homeostasis model or fasting plasma insulin level, it had a significant correlation with the fasting C-peptide level (Supplemental Fig. 1 A to C). However, it was not correlated with blood pressure or the fasting glucose level (Supplemental Fig. 1D to F). There were no difference in OCR, ATP content and the ROS content of the cybrids harboring mtDNA haplogroups A, B, D and F as a group (data not shown).

**Figure 1.**
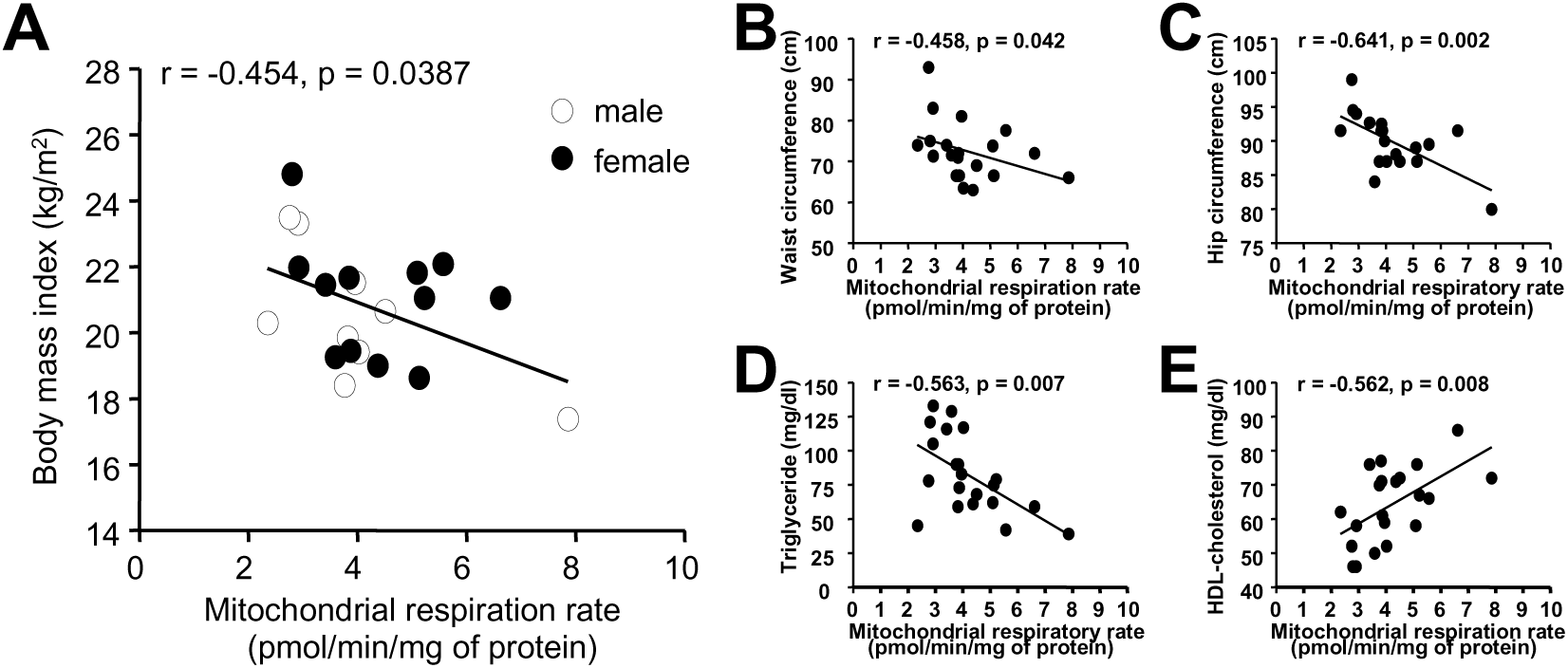
Correlations between mitochondrial oxygen consumption rates and components of the metabolic syndrome. Correlation with body mass index (A), waist circumference (B), hip circumference (C), serum triglyceride concentrations (D), and serum HDL-cholesterol concentrations (E) were shown. The mitochondrial respiration rate was calculated by subtracting the non-mitochondrial oxygen consumption rate (measured after KCN treatment) from the basal oxygen consumption rate. Each point of mitochondrial respiration is the mean of 6 independent measurements. Pearson’s correlation coefficients and P-values were shown.

### *In vitro* cell proliferation rate

We initially compared the cell numbers for 72 h after seeding the same number of cells. The proliferation rates of cybrids harboring normal mitochondrial genomes were much lower than the parental 143B TK^‒^ ρ^+^ cells (Fig. 2A). Among the mtDNA haplogroups, haplogroup D showed the lowest proliferation rate as compared to the others (*P* < 0.05 by Mann-Whitney U test) (Fig. 2B). However, we could not find any difference in the basal OCR, ATP content and the ROS content of the cybrids harboring mtDNA haplogroups A, B, D and F (Supplemental Fig2).

**Figure 2.**
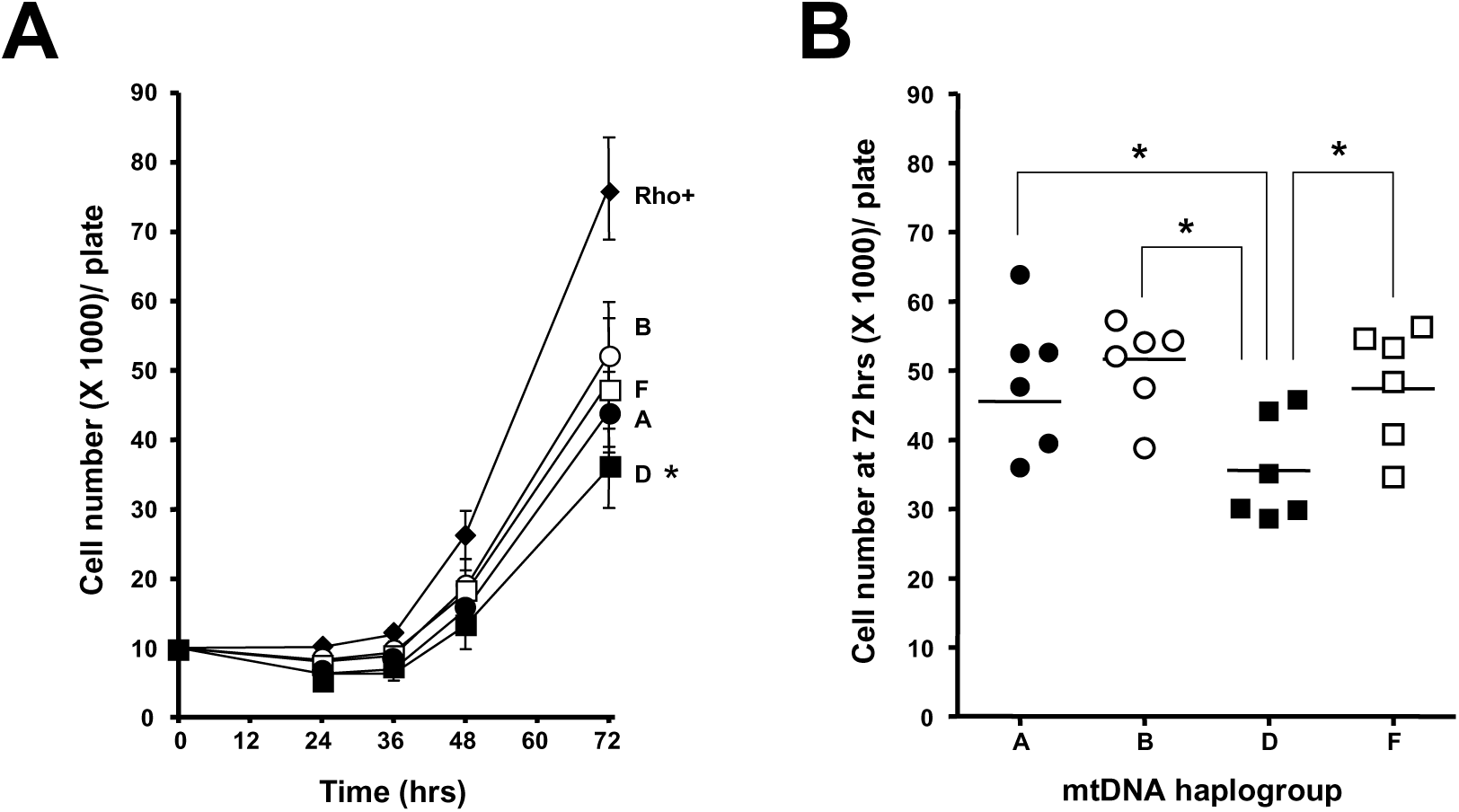
Cell proliferation rates of cybrids. (A) Cell proliferation rates of cybrids were compared to 143B TK^‒^ ρ^+^ cells, which have the same nuclear background with all cybrids. Data were shown as mean ± SD of 6 clones in each haplogroup. Cell numbers of individual cybrid clones were measured at 24, 36, 48, and 72 hrs after seeding the same number of cells (1 × 10^4^ cells) in three independent experiments. *, P < 0.05 compared to ρ^+^ cells. (B) Comparison of cell number at 72 hours among cybrids harboring each mtDNA haplogroup showed significant difference. P-values were from Mann-Whitney U test. Each point represents the mean of three independent measurement of each clone (six clones per haplogroup). *, P < 0.05.

To further examine the mechanism of the different proliferation rates, we selected representative rapid- and slow-growing cells. Among 24 cybrids, when we ranked the growth rate of each cybrid clone (data not shown), one clone harboring haplogroup A, designated as A1, had the fastest growth, and 3 cybrids harboring haplogroup D, designated as D1, D3 and D6, were the slowest to grow. The thymidine incorporation rate was higher in cybrid A1 as compared to that in cybrids D1, D3 and D6 (Fig. 3A). There was no difference in the rate of apoptosis which was examined by the terminal deoxynucleotidyl transferase (TdT)-mediated deoxyuridine triphosphate (dUTP)-biotin nick end-labeling (TUNEL) method (Fig. 3B). There was no difference in ROS production among cybrids A1, D1, D3 and D6, while 143B TK^‒^ ρ^+^ cells showed significantly increased levels of ROS, which is known as mitogenic factor (Petros et al. 2005) (Fig. 3C). Also we could not find any differences in the mitochondrial membrane potential, basal OCR and cellular ATP content among these cybrids (Fig. 3D to F).

**Figure 3.**
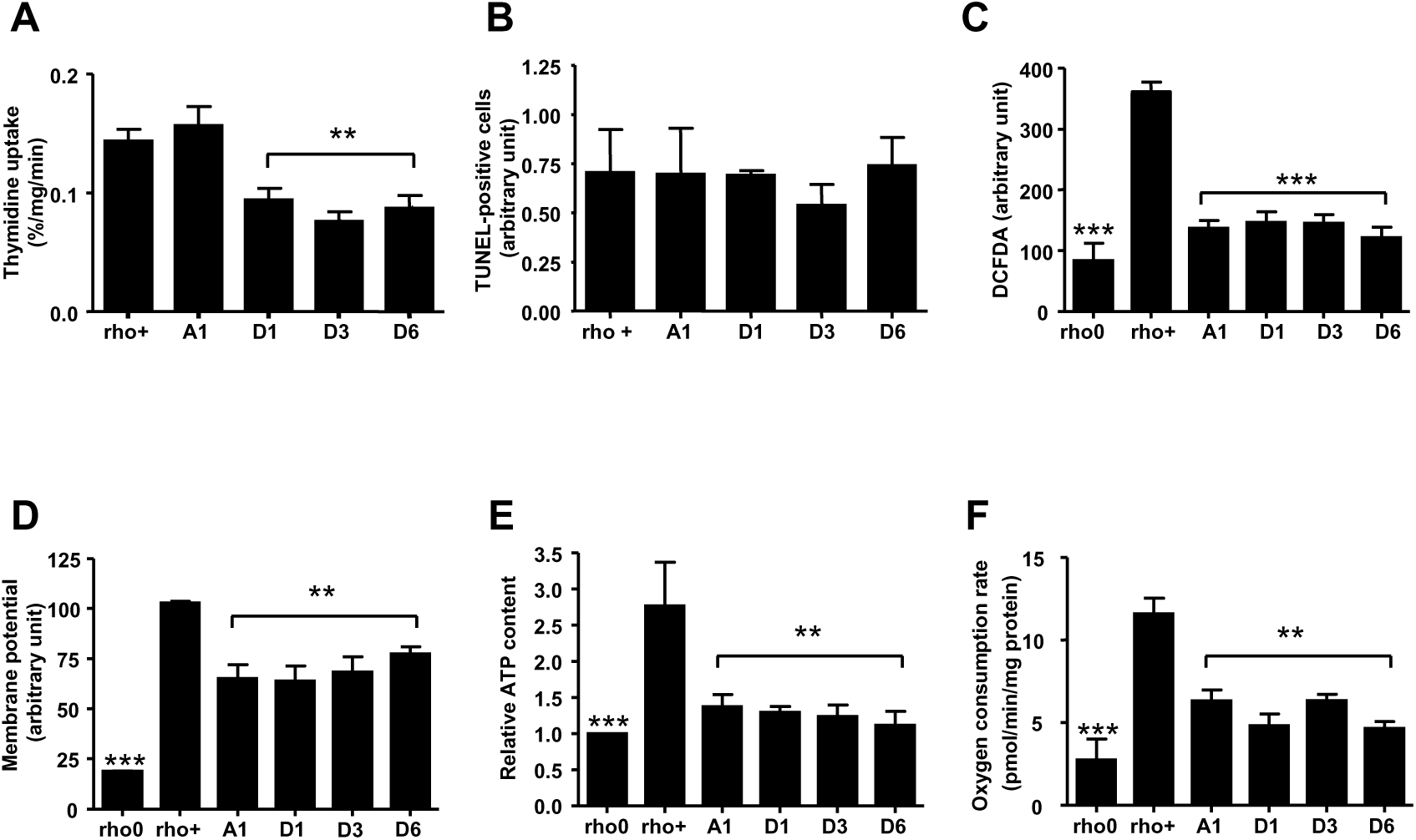
Cell functions regarding to growth and mitochondria. (A) Thymidine incorporation rates of cybrid D1, D3, and D6 were significantly lower than 143B TK^‒^ ρ^+^ (rho+) and cybrid A1 cells. **, P < 0.01 compared to rho+ and A1. (B) Apoptosis rates were comparable in all cell types. (C) Intracellular ROS contents measured by DCFDA intensity were significantly lower than rho+ cells in rho0 and cybrids A1, D1, D3, and D6, while they were comparable among cybrids. ***, P < 0.001 compared to ρ^+^. (D) Mitochondrial membrane potentials of cybrids measured by TMRE intensity were lower than rho+, while they were comparable among cybrids. **, P < 0.01; and ***, P < 0.001 compared to ρ^+^. (E) Intracellular ATP contents relative to rho0 cells were lower than ρ^+^, while they were comparable among cybrids. **, P < 0.01; and ***, P < 0.001 compared to ρ^+^. (F) Basal oxygen consumption rates were lower than ρ^+^, while they were comparable among cybrids. **, P < 0.01; and ***, P < 0. 001 compared to ρ^+^. P-values were calculated by Kruskal-Wallis test with Dunn’s multiple comparisons test.

### *In vivo* tumorigenicity and angiogenesis

In order to examine the *in vivo* tumorigenicity, we inoculated 5 × 10^6^ cells of 143B TK^‒^ ρ^+^ and cybrid clones A1, B5, D1, D3 and F1 into BALB/c athymic nude mice and observed tumor growth *in vivo* for 3 weeks (Supplemental Fig. 3). Cybrid A1 grew faster than the other cybrids and the 143B TK^‒^ ρ^+^ cells (*P* < 0.05). Further, cybrids D1 and D3 again showed the slowest growth rate. On examination under a light microscope, no massive necrotic cell death was observed. We stained tumor sections with platelet/endothelial cell adhesion molecule-1 (PECAM-1 or CD31) in order to observe the capillary density in the tumor. The capillary density in the tumor formed by 143B TK^‒^ ρ^+^ and cybrid A1 cells was significantly higher compared to those of cybrids D1 and D3 (Fig. 4A and B). To examine the *in vitro* angiogenic effect of each cybrid clone, we performed a matrigel tube formation assay using human umbilical venous endothelial cells (HUVECs) and the conditioned media from A1 and D3. The tube-forming ability of cybrid A1 was greater than that of D3 (Fig. 4C and D).

**Figure 4.**
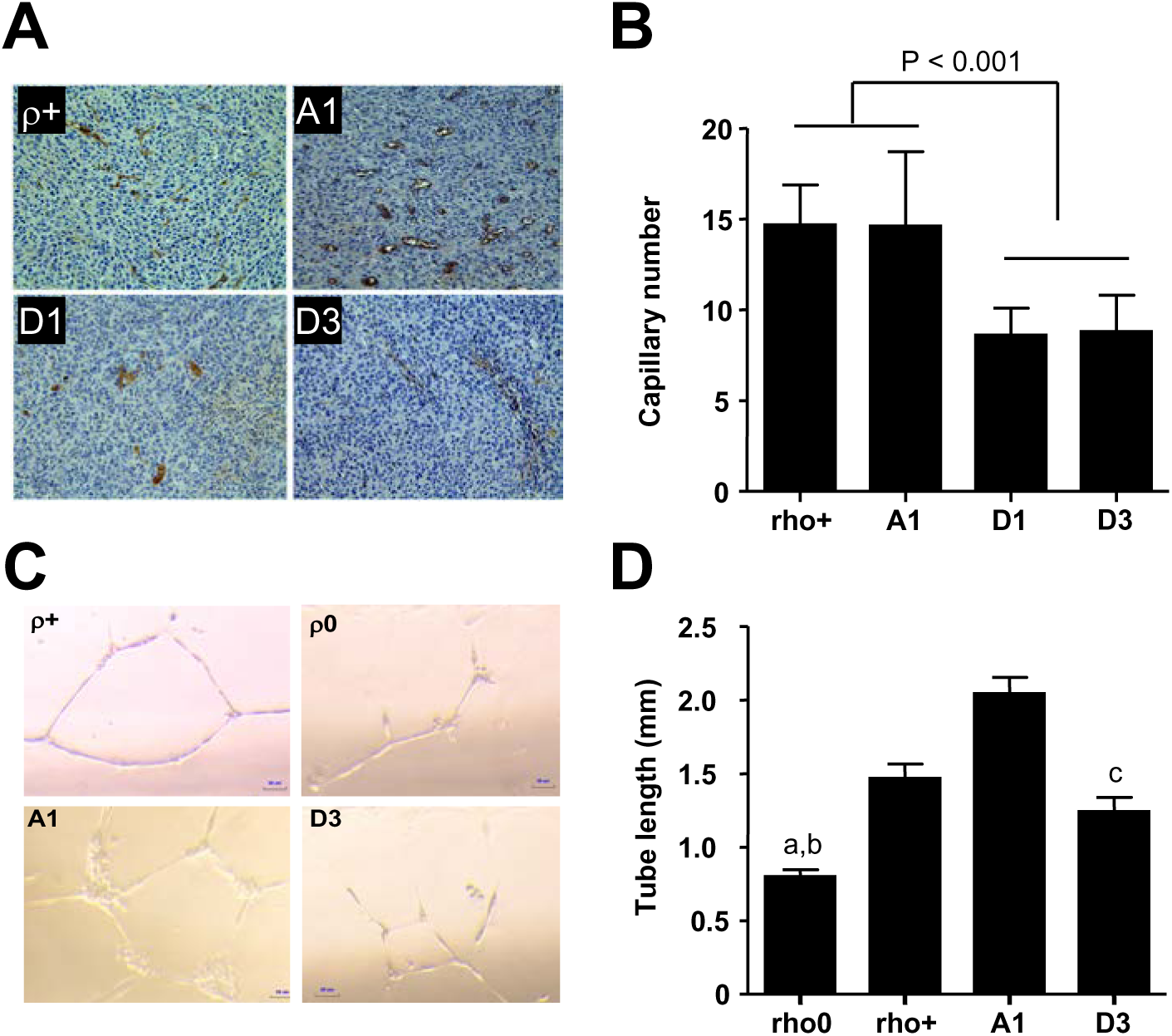
*In vivo* and *in vitro* angiogenesis. (A) Representative photographs of 21-day-old tumors of 143B TK^‒^ ρ^+^ cell and cybrids A1, D1, and D3 stained with CD31 (PECAM-1) at magnitude x 200 were shown. (B) The results of quantitative analysis of capillary density in tumors of 143B TK^‒^ ρ^+^ cell and cybrid A1, D1, and D3 were shown. Eight fields per mass were examined by 3 independent examiners blinded for the study design. Three mice for one cybrid clone were subjected for this study. Data were shown as mean ± SD of capillary number of tissue section examined at magnitude × 200. (C) HUVECs were seeded to the chamber slide with 400 μL of mixture of EGM media and conditioned media from each cybrid (1:1) and then were observed under an inverted microscopy 12 hours later. Representative photographs were shown. (D) For quantitative analysis, five representative fields were taken and the average of the total length of the tubes was compared. a, P < 0.05 compared to 143B TK^‒^ ρ^+^; b, P < 0.001 compared to A1; c, P < 0.05 compared to A1 by Kruskal-Wallace test and Dunn’s multiple comparisons test (n = 8)

### Comparisons of mtDNA sequences and differential gene expression

Cybrid A1 was the fastest growing cell both *in vitro* and *in vivo* among the cybrids with normal mtDNA. By sequencing whole mtDNA (Supplemental Tables 1 to 26), we found that cybrid A1 had a T7372C (Met490Thr) mutation in the cytochrome *c* oxidase I (*COI*) gene and an A12715G (Thr127Ala) mutation in the NADH dehydrogenase subunit 5 (*ND5*) gene (Supplemental Table 1).

We hypothesized that the differences in the tumorigenicity and cell proliferation rate might be explained by differential nuclear gene expression in response to the yet unidentified changes originated from different mitochondrial genome. Using cDNA microarrays, we compared the gene expression profiles of cybrids A1 and D3, i.e., the cells with the fastest and slowest growth, respectively. In cybrid A1, the expression levels of 23 genes were increased (Supplemental Table 27) and those of 11 genes were decreased (Supplemental Table 28) as compared to the expression levels in cybrid D3. Among these genes, thrombospondin-1 (*THBS1*) expression was markedly increased in cybrid A1 (9.14 fold), and it was confirmed by northern blot analysis (Supplemental Fig. 4A). Several genes increased in expression level in cybrid A1 as compared to D3 (asparagine synthetase [ASMS], fibroblast growth factor 13 [*FGF13*], insulin-like growth factor binding protein 3 [*IGFBP3*], RAB4A, member RAS oncogene family [*RAB4A*] and *THBS1*); this was confirmed by RT-PCR (Supplemental Fig. 4B).

## Discussion

Mitochondrial dysfunction is known to be a major contributor to the pathogenesis of metabolic syndrome or type 2 diabetes. A decreased capacity of mitochondrial OXPHOS has been shown to be associated with insulin resistance, which is a key feature of the metabolic syndrome and type 2 diabetes, in aged people and in the offspring of type 2 diabetic patients (Petersen et al. 2003; Petersen et al. 2004). Microarray studies have also shown that insulin resistance is associated with the decreased expression of genes related to OXPHOS in skeletal muscles (Mootha et al. 2003; Patti et al. 2003). A control region polymorphism T16189C is known to be associated with insulin resistance, obesity and diabetes in both Europeans (Poulton et al. 2002) and Asians (Kim et al. 2002; Weng et al. 2005). A new homoplasmic mutation substituting cytidine for uridine immediately 5′ to the mitochondrial transfer RNA(Ile) anticodon was found to be the cause for the familial clustering of metabolic disorders such as hypertension, hypercholesterolemia and hypomagnesaemia (Wilson et al. 2004). Further, we have recently found that a common mtDNA haplogroup N9a is associated with the resistance to develop type 2 diabetes both in Japanese and Koreans (Fuku et al. 2007). In this study, the mitochondrial OCR of cybrids, the differences of which were solely determined by the differences in mitochondrial genome, was associated with body mass index, waist circumference, triglyceride, and HDL-cholesterol levels. Although we did not obtain a significant correlation between mitochondrial OCR and insulin resistance assessed by the homeostasis model, there was a modest but significant correlation between the mitochondrial OCR and fasting C-peptide levels, which was known to be associated with insulin resistance (Chen et al. 1999). Interestingly, it is noteworthy that these statistically significant correlations between mitochondrial OCR and phenotypes of metabolic syndrome were observed in the healthy subjects who were free from metabolic syndrome or diabetes mellitus. These findings along with the results from the study on conplastic rats (Pravenec et al. 2007) might add strength to the evidence addressing that the mitochondrial genomes harboring different mtDNA polymorphisms play an important role in modulating the susceptibility to the metabolic syndrome.

In the second part of this study, we examined whether mitochondrial genome might contribute to the tumorigenesis. Mitochondrial dysfunction and mtDNA mutations are linked to cancer and tumorigenesis (Brandon et al. 2006). Although there are numerous reports regarding tumor-specific somatic mtDNA mutations (Brandon et al. 2006), few examples are known on oncogenic germline mtDNA mutations. The germline 10398A allele in the *ND3* gene was linked to an increased risk of invasive breast cancer in African-American women, while the same polymorphism did not increase the breast cancer risk in Caucasian women (Canter et al. 2005). In addition, a common germline polymorphism T16189C in the hypervariable region has been shown to be associated with endometrial cancer (Liu et al. 2003). In the current study, we showed that common and apparently normal germline mtDNA polymorphisms known as haplogroup D might contribute to a protective effect against cancer by reducing tumorigenicity, which could support the fact that the mtDNA haplogroup D is associated with longevity in Japanese (Tanaka et al. 1998).

Interestingly, the cell proliferation rate and tumorigenicity of the 143B TK^‒^ ρ^+^ cells were dramatically reduced when their mitochondrial genome was substituted with that of normal individual with exception of cybrid A1. By sequencing whole mtDNA, we found that cybrid A1 had a T7372C (Met490Thr) mutation in the *COI* gene and an A12715G (Thr127Ala) mutation in the *ND5* gene. T7372C is a new mutation, and 127Thr (A at nucleotide position 12715) is a highly conserved amino acid among 61 mammalian species (http://mtsnp.tmig.or.jp/mtsnp/index_e.shtml). Therefore, these variants might be responsible for the functional differences of cybrid A1, although the functional significance of each variant at the molecular level is still elusive.

It has been reported that increased levels of ROS play an important role in the tumorigenesis of prostate cancer cells (Petros et al. 2005). We also found that 143B TK^‒^ ρ^+^ cells produced much more ROS than cybrids with normal mitochondrial genome. However, there was no significant difference in the ROS levels between A1 and other cybrids, the increased tumorigenicity of cybrid A1 must be related to unknown mitogenic factors other than ROS. Of note, many nuclear DNA-encoded genes showed various expression levels depending on the variations in mitochondrial genome in spite of the same nuclear background.

In conclusion, we showed that the mitochondrial genome from healthy individuals might play a role in the pathogenesis of common complex diseases in humans. These findings suggest that the effect of the individual mitochondrial genome as well as that of the nuclear genome should be considered in explaining the genetic pathogenesis of common complex human diseases.

## Methods

### Human subjects and clinical measurements

The Institutional Review Board of the Clinical Research Institute in Seoul National University Hospital approved the study protocol, and an informed consent was obtained from each subject for the genetic analysis. We enrolled a total of 24 subjects harboring the common Asian mtDNA haplogroups A, B, D and F (n = 6 for each haplogroup). There were 11 men and 13 women (age, 40–63 years). At the time of the study, all the subjects were free from any form of clinically diagnosed cancer. Among them, 21 subjects (9 men, 12 women; age, 40‒63 years) did not have diabetes, obesity or metabolic syndrome, i.e., that might cause secondary mitochondrial dysfunction (Kelley et al. 2002). The mtDNA haplogroups were initially screened by the PCR-restriction fragment length polymorphism (RFLP) method as described elsewhere (Wallace et al. 1999) and were confirmed by the direct sequencing of whole mtDNA by methods described previously (Kazuno et al. 2006). All the sequence variations are shown in the supplemental tables 1‒26. All the study subjects were examined in the morning after an overnight fast. The height, weight, circumferences of waist and hip and blood pressure were measured. Blood samples were drawn for biochemical measurements (fasting plasma glucose, fasting plasma insulin, C-peptide, HbA1c, total cholesterol, triglyceride and HDL-cholesterol) and DNA extraction.

### Cell culture and platelet-mediated transformation of human mitochondrial DNA-less cells

The osteosarcoma cell line with neither mtDNA nor thymidine kinase activity (143B TK^‒^ ρ^0^), which was obtained after long-term exposure to ethidium bromide (50 ng/ml), was grown in Dulbecco’s modified Eagle’s medium (DMEM; GibcoBRL) supplemented with 100 mg/ml 5-bromodeoxyuridine (BrdU), 50 μg/ml uridine, and 10% fetal bovine serum (FBS). Southern blot analysis and PCR amplification of the mtDNA target sequences confirmed the absence of any residual mtDNA. Using platelets as mtDNA donors, cybrids were produced as described previously (Chomyn 1996). To assure a complete repopulation of mtDNA, the functional assessment of the selected clones was carried out after 2‒3 months of successive subcultivation.

### Measurement of oxygen consumption rates

Cellular oxygen consumption rates (OCRs) were measured as previously described with a slight modification (Villani and Attardi 2001; Yoon et al. 2005). Briefly, the exponentially growing cells (5 × 10^6^) were cultured in DMEM supplemented with 10% FBS, washed with PBS buffer (pH 7.2) and collected by trypsinization. After resuspending the cells in 1 ml complete medium without phenol red, the cells were transferred to the Mitocell chamber equipped with a Clark-type oxygen electrode (Strathkelvin). OCR was measured at the baseline and in the presence of 30 μM 2,4-dinitrophenol or 100 μM KCN for uncoupled respiration and non-mitochondrial respiration, respectively.

### Measurement of intracellular ATP content

The ATP in the cell was measured using the luciferin-luciferase reaction with an ATP bioluminescence somatic cell assay kit (Sigma) according to the manufacturer’s instructions. The luminescence was measured by a bioluminometer (Berthold). The ATP content was adjusted for the amount of protein that was measured by the Bradford method.

### Measurement of intracellular ROS content

The intracellular ROS content was measured by using 2′,7′-dichlorofluorescein diacetate (DCFDA; Invitrogen). The cells were plated on 6-well plates and incubated with 1.5 μl/ml DCFDA at 37°C for 30 min. ROS in the cells cause the oxidation of DCFDA, yielding the fluorescent product 2′,7′-dichlorofluorescein (DCF). The DCF fluorescence was measured in intact cells by fluorescence-activated cell sorting analysis using FACSCalibur (Becton Dickinson).

### *In vitro* cell proliferation rate measurement

We measured the cell numbers using a hemocytometer at 24, 36, 48 and 72 h after seeding the same number of cells (1 × 10^4^ cells/100 mm dish).

### [^3^H]-thymidine incorporation rate measurements

A [^3^H]-thymidine incorporation assay was used to evaluate cell proliferation. Each cybrid was seeded in a 24-well plate. At the 80% confluent state, the cells were made quiescent by incubation in DMEM without FBS for 24 h and were then incubated in DMEM containing 10% FBS. Simultaneously, 1 μCi/ml of [^3^H]-thymidine was added to the media and incubated for 4 h. The cells were washed with PBS, fixed in cold 10% trichloroacetic acid, washed with 95% ethanol and dried. The incorporated [^3^H]-thymidine was extracted in 0.2 M NaOH and measured in a liquid scintillation counter. The [^3^H]-Thymidine incorporation results were normalized by a negative control value, and the protein amount was measured by the Bradford method and expressed as %/min/mg.

### Apoptosis assay

Apoptosis of the cybrid at 80% confluence was detected by immunofluorescence staining in a species-independent TUNEL assay according to the protocols of the supplier (APO-BrdU TUNEL assay kit; Molecular Probes). Briefly, DNA 3′-hydroxyl ends exposed by nucleases as a consequence of apoptosis were labeled with 5-bromo-2′-deoxyuridine (BrdU) 5′-triphosphate (BrdUTP) by TdT. The BrdU incorporated into the DNA was detected with the Alexa Fluor 488-conjugated anti-BrdU mouse monoclonal antibody PRB-1 (Molecular Probes) by flow cytometry using FACSCalibur (Becton Dickinson).

### Transplantation of cybrids to form tumors in nude mice

Cybrid cells suspended in Matrigel (5 × 10^6^ cells/0.2 mL) were injected subcutaneously into the scapular area of male athymic mice (4-week-old BALB/c *nu*/*nu*). For 3 weeks after transplantation, the tumor volume was measured at 3–4 day intervals using the formula *V* = (*A* × *B*^2^)/2, where *V* is the volume (mm^3^), *A* is the long diameter (mm), and *B* is the short diameter (mm). At 3 weeks after transplantation, the mice were sacrificed and the tumors were removed for further analysis.

### Tissue histochemistry and quantification of capillary density

The tumors were removed 21 days after transplantation and fixed overnight in 10% formalin at 4°C. The fixed tissues were embedded in paraffin and sectioned at 5 μm. The capillary density was examined by the quantification of endothelial cells in cryostat sections stained with antimouse CD31 (PECAM-1; Santa Cruz), an antibody specific for mouse endothelial cells, following standard immunoperoxidase procedures. Eight non-overlapping microscopic fields at a magnification of ×200 were analyzed from each tumor by 3 persons who were blinded to the study protocol.

### In vitro capillary formation

A Matrigel (Becton Dickinson) basement membrane matrix was thawed and mixed in the ratio of 1:1 with endothelial growth media (EGM)-2 (Cambrex). This mixed Matrigel (100 μm) was prelayered onto a chamber slide. After 1 h incubation at room temperature, 1 × 10^4^ HUVECs were seeded onto the chamber slide with a 400-μm mixture of EGM media and conditioned media from each cybrid (1:1). The slide was observed under an inverted microscope after 12 h. Five representative fields were taken, and the average of the total length of the tubes was compared.

### Microarray analysis

The total RNA from the cells was prepared using the RNeasy midi kit (Qiagen). The Applied Biosystems Human Genome Survey Microarray contains 33,096 60-mer oligonucleotide probes representing 27,858 individual human genes. Digoxigenin-UTP labeled cRNA was generated and amplified from 2 μg of total RNA from each cybrid at 80% confluence using Applied Biosystems Chemiluminescent RT-IVT Labeling Kit v 2.0 according to the manufacturer’s protocol. Array hybridization was performed for 16 h at 55°C. Chemiluminescence detection, image acquisition and analysis were performed using the Applied Biosystems Chemiluminescence Detection Kit and Applied Biosystems 1700 Chemiluminescent Microarray Analyzer according to the manufacturer’s protocol. The images were auto-gridded and the chemiluminescent signals were quantified, background subtracted, and finally, spot and spatially normalized using the Applied Biosystems 1700 Chemiluminescent Microarray Analyzer software v 1.1.1.

### RT-PCR

The total RNA from the cells was prepared using Trizol (Invitrogen). cDNA was obtained using 1 μg of RNA with random hexamers and avian myeloblastosis virus reverse transcriptase (Invitrogen). The PCR reactions were carried out with 5 μL of cDNA template, 10 pM of each primer, 8 μL of 2.5 mM dNTP mix and 0.1 units of rTaq DNA polymerase (Takara Bio) in a volume of 50 μL. The primer sequences for RT-PCR are shown in Supplemental Table 29. The samples were amplified in a thermocycler under the following conditions for 17–25 cycles: first, the denaturing step at 95°C for 30 s, then the annealing step at 50–60°C for 40 s and finally the amplification step at 72°C for 40 s.

### Statistical analysis

Data are shown as means ± SD. For the comparison of cellular or mitochondrial function, the Mann-Whitney U test, Kruskal-Wallis test or Pearson’s correlation analysis were applied where appropriate. All the tests were performed with a commercially available statistical package (SPSS).

## Acknowledgments

This work was supported by grants from the Korea Health 21 R & D Project, Ministry of Health & Welfare, Republic of Korea (02-PJ1-PG1-CH04-0001 to H.K.L. and 00-PJ3-PG6-GN07-001 to K.S.P. and Y.M.C.), MIC & IITA through IT-Leading R&D Support Project (to H. K.L.) and FPR05C2-450 of 21C Frontier Functional Proteomics Project from the Korean Ministry of Science & Technology (to Y.K.P.). We thank Dr. Yau-Huei Wei for providing 143B TK^‒^ ρ^0^ cells and Ji Hyeon Kim and Hae Sun Kang for their excellent technical assistance.

## Web site references

http://mtsnp.tmig.or.jp/mtsnp/index_e.shtml

